# Mediodorsal thalamus is critical for updating during extra-dimensional shifts but not reversals in the attentional set-shifting task

**DOI:** 10.1101/2021.04.13.439610

**Authors:** Zakaria Ouhaz, Brook AL Perry, Kouichi Nakamura, Anna S Mitchell

**Author notes:** Correspondence should be addressed to ASM. Author contributions: ZO and ASM designed research; ZO performed research; KN and BALP contributed unpublished reagents/ analytic tools; ZO, BALP, ASM analyzed data; ZO, BALP, KN, ASM wrote the paper.

## Abstract

Cognitive flexibility, attributed to frontal cortex, is vital for navigating the complexities of everyday life. The mediodorsal thalamus (MD), interconnected to frontal cortex, may influence cognitive flexibility. Here male rats performed an attentional set-shifting task measuring intra-dimensional and extra-dimensional shifts in sensory discriminations. MD lesion rats needed more trials to learn the rewarded sensory dimension. However, once the choice response strategy was established, learning further two-choice discriminations in the same sensory dimension, and reversals of the reward contingencies in the same dimension, were unimpaired. Critically though, MD lesion rats were impaired during the extra-dimensional shift, when they must rapidly update the optimal choice response strategy. Behavioral analyses showed MD lesion rats had significantly reduced correct within trial second choice responses. This evidence shows transfer of information via the MD is critical when monitoring and rapid within trial updates in established choice response strategies are required after there is a rule change.

**Significance statement:** We demonstrate for the first time that rodent mediodorsal (MD) thalamus is a critical node when choice response strategies need to change rapidly after a within session rule change but not after reversals of reward contingencies during reward guided learning. MD interactions with orbitofrontal cortex are critical for value based learning, while MD interactions with medial prefrontal cortex are critical for rapid within trial updating of optimal choice response rules. MD interactions with the orbitofrontal cortex are not always necessary for reversal learning.

## Introduction

Cognitive flexibility describes our ability to quickly and selectively switch our thoughts, responses, or behavior to everyday dynamic situations. This capacity to rapidly update or alter one’s actions conveys evolutionary benefit and is key to survival (Koechlin et al., 2003; Rougier et al., 2005). Cognitive flexibility shows marked changes or declines in neurodegenerative disorders, like Parkinson’s disease, and in neurodevelopmental diseases, like schizophrenia (Andrews et al., 2006; Barch et al., 2009; Giraldo-Chica et al., 2018).

Cognitive flexibility is not unique to primates. Studies show that rats also can readily switch between attentional sets to optimise reward outcome (Birrell and Brown, 2000; McAlonan and Brown, 2003; Newman and McGaughy, 2011). Typically, these studies have focussed on various frontal cortex subregions. Yet, it is becoming increasingly clear that the cortex does not function in isolation, but rather, relies heavily on subcortical and peripheral inputs supplied by the thalamus (Sherman and Guillery, 2002; Guillery and Sherman, 2011; Halassa and Kastner, 2017). Clearly, the mediodorsal thalamus (MD) has a critical role in functions of frontal cortex during higher order cognitive processes across mammalian species (Alcaraz et al., 2018; Browning et al., 2015; Chakraborty et al., 2016; DeNicola et al., 2020; Ferguson and Gao, 2018; Miller et al., 2017; Pergola et al., 2018; Parnaudeau et al., 2013; Perry et al., 2021; Schmitt et al., 2017). Monkeys with MD lesions were found to have deficits when required to rapidly learn and update trial-relevant information during complex visuospatial associative tasks (Browning et al., 2015; Chakraborty et al., 2016; Mitchell et al., 2007a) but not during recognition of preoperative acquired visuospatial discriminations (Mitchell and Gaffan, 2008), or implementation of a preoperatively learnt response strategy (Mitchell et al., 2007a). This evidence highlights that the MD and PFC are working in partnership, contributing different but complementary roles to goal-directed behaviours (Mitchell, 2015; Perry et al., 2021). In monkeys and humans, the MD is densely interconnected with frontal cortex areas involved in various aspects of cognitive flexibility, namely the dorsolateral and dorsomedial prefrontal cortex (PFC), equivalent to the medial PFC (prelimbic (PrL)) in rodents, the orbitofrontal cortex (OFC), ventrolateral PFC, and the dorsal anterior cingulate cortex (Dajani and Uddin, 2015; Perry et al., 2021; Preuss, 1995). Recent neuroanatomical work in rodents has shown the arrangement of MD projections to the frontal cortex allow for the simultaneous sharing of information across multiple PFC cortical regions (Alcaraz et al., 2016; Kuramoto et al., 2017). This distributed pattern of prefrontal innervation by the MD appears to be especially true for the central (MDc) and medial (MDm) subdivisions, which have projections to OFC, medial PFC, and frontal association areas (Alcaraz et al., 2016; Groenwegen, 1988). Although phylogenetic differences exist in the organisation of rodent and primate frontal cortex, similar functional contributions to aspects of cognitive flexibility are observed in monkeys and rodents after perturbations to similar areas of agranular frontal cortex (Birrell and Brown, 2000; Bissonette et al., 2008; Dias et al., 1996; Dias et al., 1997; McAlonan and Brown, 2003). Given there are similarities in neuroanatomical connectivity between agranular frontal cortex and MD in rodents and primates, it seems that cognitive flexibility and rapid updating deficits observed in monkeys with MD lesions may also extend to rodents.

Thus, in the current study, rats with excitotoxic lesions to the MD or MD Sham controls were run on a well-established test of cognitive flexibility – the intra-dimensional (ID)/extra-dimensional (ED) attentional set-shifting task (Birrell and Brown, 2000; Tait et al., 2014; Tait et al., 2018) to test this premise and investigate any deficits in choice responding. We used the standard 7-stage version comprising of multiple subtasks all conducted within a single testing session (Birrell and Brown, 2000; Chase et al., 2012). In rats, the ID/ED task has produced dissociable deficits after permanent cortical or thalamic lesions, or neuropharmacological manipulations (e.g., Linley et al., 2016; McGaughy et al., 2008; Tait et al., 2014; Tait et al., 2018; Wright et al., 2015).

In the current experiment, we predicted MD lesion rats would show deficits during the initial sensory acquisition (simple discrimination, SD) and the ED shift. Both the SD and ED subtasks require rapid, within trial learning or updating of choice response strategies. Therefore, impaired SD and ED performance would be consistent with selective deficits in learning observed in other rodent studies after MD perturbations (Chakraborty and Mitchell, 2013, for review; Courtiol et al., 2019; Ferguson et al., 2017; Hunt and Aggleton, 1998; Mitchell and Dalrymple-Alford, 2005; Perry et al., 2021). We also predicted our MD lesion rats may show deficits during reversal subtasks given the reciprocal direct connectivity with the OFC (Groenwegen, 1988; Krettek and Price, 1977; Ray and Price, 1992).

## 2.0 Materials and Methods

### Animals

Twenty-five Lister hooded male rats weighing between 420 and 480 gm at time of surgery (Charles River, UK) were group housed in a temperature and humidity-controlled environment (21 ± 1°C). The housing and husbandry compiled with the ARRIVE guidelines of the European Directive (2010/63/EU) for the care and use of laboratory animals. Testing was conducted in the light phase of a 12hr light/dark cycle (lights on at 7am). The rats were maintained on a controlled feeding schedule (20 g/rat/day) with water freely available in the home cage. All experimental procedures were performed in compliance with the United Kingdom Animals (Scientific Procedures) Act of 1986. A project license authorized all procedures after review by the university animal care and ethical review committee and Home Office Inspectorate.

### Surgery

Rats were anesthetized by isoflurane and oxygen mix (4% induction, 1.8–2% maintenance), and given analgesia in subcutaneous injections of 0.05mg/kg buprenorphine (Vetergesic; 0.3 mg/ml) and 1 mg/kg Metacam (Meloxicam; 5 mg/ml). Rats were secured in a stereotaxic frame (Kopf) with atraumatic ear bars and the nose bar set to +3.3 mm to achieve a level skull. A subcutaneous injection of 2 mg/kg bupivacaine (Marcaine; 2.5 mg/ml) was administered into the scalp in the location of the midline incision. Viscotears was applied to keep the eyes moist. During surgery, each rat was placed on a heat pad and covered in bubble wrap with an internal rectal thermometer probe to monitor and maintain normal body temperature. Warmed sterile saline (1 ml/100 gm) was administered subcutaneous into the scruff of the neck after one hour. A midline incision was performed. Bregma and lambda coordinates were determined. A dental drill with a trephine head was used to create a craniotomy over the midline. To maximize lesion accuracy, anterior and posterior injection sites were calculated according to the bregma–lambda distance in each rat. Coordinates were for anterior MD injection: anterior–posterior (AP) = – 0.395 mm, medial-lateral (ML) = – 0.1 (avoiding the superior sagittal sinus, which was visible within the dura inside the craniotomy) and dorsal-ventral (DV) = 0.56 mm (from dura), volume of excitotoxin = 0.20 µl; for posterior MD: AP = – 0.435 mm, ML = + 0.1 and DV = 0.57 mm (from dura), volume of excitotoxin = 0.18 µl. Fifteen rats (MD lesion) received 0.12M of NMDA (N-methyl-D-Aspartate) dissolved in phosphate buffer (pH 7.20) in each hemisphere from a 1 µl-gauge Hamilton bevelled-tip syringe. The injections were performed manually, taking 3 min each, and after injection the needle was left in situ for a further 3 min to allow diffusion and to limit wicking of the NMDA. A further 10 rats (MD Sham controls) received injections of sterile phosphate buffer instead of NMDA using the same injection coordinates. Injections were given bilaterally in the MD. Upon completion of surgery, wounds were sutured with Vicryl 4.0. Rats were housed singly during recovery for up to three days and were then returned to their pre-surgery housing cohort. Postoperative oral analgesia, 1mg/kg Metacam (Meloxicam, 1.5 mg/ml) dissolved and set in jelly was provided in individual bowls to each rat daily for three days. Behavioral and physiological evidence showed that all rats recovered well, with normal eating and pre-surgery weights returning within 24h. Food regulation (20 gm/day/rat) started again 10d after recovery. Postoperative testing began at least 15d after surgery. The researcher was blind to the lesion group of each animal until all tests were completed.

### ID/ED Attentional Set Shifting paradigm

#### Apparatus

The test chamber consisted of a modified Plexiglass home cage (40 × 70 × 18 cm) with Perpex dividers separating the cage into a large start chamber (40 × 46 cm) and two identically sized (24 × 20 cm) choice chambers to which access was controlled by removable Perspex doors. The digging bowls were ceramic (internal diameter 7 cm, depth 4 cm) and were placed within the choice chambers. The bowls were filled with digging media of different textures, and the digging media were scented with different spices (see **Table 1** for the examples used). The odor or digging media discriminations, pairs of stimuli used, and the correct stimulus within pairs were counter-balanced across subtasks and matched between groups. The bowls were baited with Honey Nut Cheerios (Nestle®), each bowl contained a hidden (securely fixed under a metal grid) Honey Nut Cheerio to serve as an odor mask. During testing, only one of the bowls was baited and rats determined which bowl was baited using either the texture of the digging medium or the odor as cues. Prior to testing, rats were taught to dig in bowls filled with home-cage bedding material to retrieve one half of a Honey Nut Cheerio. The task was performed as previously described in Birrell and Brown (2000) and divided into two phases administered on two consecutive days.

#### Preoperative and Postoperative Training Day

On the day before testing, each rat learnt one simple two-choice discrimination (SD) using either of the two sensory dimensions: odor (mint vs oregano), or digging media (shredded paper vs cotton pads), to a criterion of six consecutive correct trials. The rewarded odors or digging media were counterbalanced across the two groups, and these exemplars were not used again throughout testing. Digging was defined as active digging with both front paws or active foraging with the snout in the digging media. Sniffing or touching the media with the paws was not scored as a dig.

#### Preoperative and Postoperative Testing paradigm

The following day, each rat was given a series of seven two-choice discriminations (subtasks) in the following order: a new simple discrimination (SD) using either of two sensory dimension (odor or digging media); a compound discrimination (CD) using the same rewarded sensory dimension and two-choice discrimination as the SD, combined with the other irrelevant sensory dimension; first reversal (REV1), in which the exemplars remained the same as in the CD but the correct (rewarded) and incorrect exemplars were reversed; intra-dimensional shift (ID), in which new exemplars were used, but the relevant sensory dimension remained the same as in the previous subtasks; second reversal (REV2), where exemplars in the ID remained the same but the reward contingencies of the two exemplars were reversed; extra-dimensional shift (ED), where new exemplars were used and the previously irrelevant sensory dimension becomes relevant; and a third reversal (REV3), where exemplars in the ED subtask remained the same but the reward contingencies of the two exemplars were reversed.

The task is self-paced and relies on the natural foraging behaviors of the rats. The first five two-choice discrimination subtasks (SD, CD, REV1, ID, REV2) must be solved by discriminating between exemplars from the same sensory dimension. In this stage of the task, the rat is rewarded for choices based on specific perceptual features of the stimuli, while ignoring other features that also distinguish the stimuli. After acquiring each set of two-choice discriminations to a performance criterion of six correct consecutive trials, the rats encounter a reversal of the reward contingencies associated with the exemplars (a reversal subtask: REV1 and REV2). For the final two subtasks (ED and REV3), exemplars from the previously irrelevant sensory dimension become relevant. Therefore the rat must shift its attention (and adjust its choice response strategy) to the previously irrelevant sensory dimension and perceptual feature to receive reward (ED subtask). The attentional set-shfiting cost is calculated by comparing trials to criterion during the intra-dimensional shift, where the sensory dimension is consistent with previous subtasks, and the extra-dimensional shift, where the now relevant sensory dimension had been previously ignored and thus requires a shift to optimise continuing to retrieve rewards.

#### Odor detection test

At the completion of ID/ ED testing, the olfactory abilities of the rat to smell the odor of the buried half-Honey Nut Cheerio were assessed. This task consisted of 10 consecutive trials where the rat was exposed to bowls containing similar bedding. The rat was placed in the waiting area of the set-shifting apparatus. Similar bedding-filled bowls were placed, one in each of the choice chambers, only one bowl contained half a Honey Nut Cheerio at the bottom (pseudorandomly assigned to avoid the rat developing a response strategy). The barrier was raised, allowing the rat access to both bowls. The rat had to chose to dig into one bowl and had up to 15 minutes to make a digging response on each trial. If the rat dug in the correct bowl, the trial was recorded as correct and the rat was returned to the waiting area and the barrier lowered to block access to both choice chambers. If the rat dug in the incorrect bowl, the trial was marked as incorrect. The rat was permitted to continue to explore the incorrect bowl; the trial was terminated when the rat returned to the waiting area. In all cases, the ability of each rat to detect the reward location was no different to chance guessing.

#### Perfusion and histology

At the end of all experimental testing, rats were deeply anaesthetized with sodium pentobarbital (Euthanol 1.0 ml/rat, 200 mg/ml: Merial, Harlow, UK), and perfused transcardially with 0.9% (w/v) saline and 4% (w/v) paraformaldehyde in 0.1M phosphate buffered saline (PBS). Brains were then removed and post-fixed overnight (4°C), then incubated using 30% (w/v) sucrose in 0.1M PBS over 48h (4°C). Free floating sections (40 µm) were cut on a freezing microtome, the slices were then preserved in cryoprotectant (0.1M PBS containing 24% (v/v) glycerol and 24% (v/v) ethylene glycol) and stored at −20°C until required. MD thalamus lesion locations were histologically verified from cresyl violet stained brain sections.

#### Lesion extent

Photomicrographs of the Nissl-stained MD sections were captured using a camera mounted on a freestanding Leica DMR microscope (Leica Microsystems) using a 1.6× objective so that the whole section could be visualised in the photomicrograph. MD volumes were measured using the NIH software ImageJ (http://rsbweb.nih.gov/ij/). In each rat, the intact MD volume from both the left and the right hemisphere was assessed. The final reading was calculated according to the Cavalieri principle (Regeur and Pakkenberg, 1989) and expressed as the percentage of MD Sham controls.

#### Behavioral data analysis

In the behavioral experiments, all rats performed a preoperative and postoperative session, and they completed the seven two-choice discrimination subtasks in an identical order. Data are expressed as mean and standard deviation. Preoperative and postoperative mean trials to criterion, shift cost, percent correct 2^nd^ choice within trial responses, and latency to dig (seconds) were analysed using repeated measures ANOVAs with Stage (comprising the 7 subtasks) as the within-group repeated measure and Lesion_type (MD lesion or MD Sham control) as the between-group factor. For significant interactions, posthoc simple main effects analyses were performed using additional two-way ANOVAs for repeated measures (e.g., ID/ED shift) or independent t-tests for relevant subtasks (corrected for multiple comparisons). Univariate ANOVA was used to determine if the Shift between the two sensory dimensions between the MD lesion or MD Sham control had an effect on the number of errors performed in the ED subtask. All statistical analyses were calculated using SPSS 24 software. The significance level was set at *p* <0.05.

## 3.0 Results

### 3.1 MD lesion site

Of the 25 adult rats involved in this experiment, 15 sustained MD lesions and 10 were MD Sham controls (see Materials and Methods for details). All 15 MD lesion rats had extensive excitotoxic (NMDA) lesion damage (see **Fig. 1**) within the medial and central subdivisions of the MD, as intended. In all cases the lesion affected between 80–90% of the central and medial MD, sparing only the lateral portions. The MD was consistently shrunken, and at the lesion site there was significant cell loss. In all cases, the adjacent anterior thalamic nuclei (ATN) were spared. In two cases, there was unilateral (right-sided) damage in the lateral habenulae (LHb) and some damage to the central medial nucleus lying underneath the MD.

**Figure 1.**
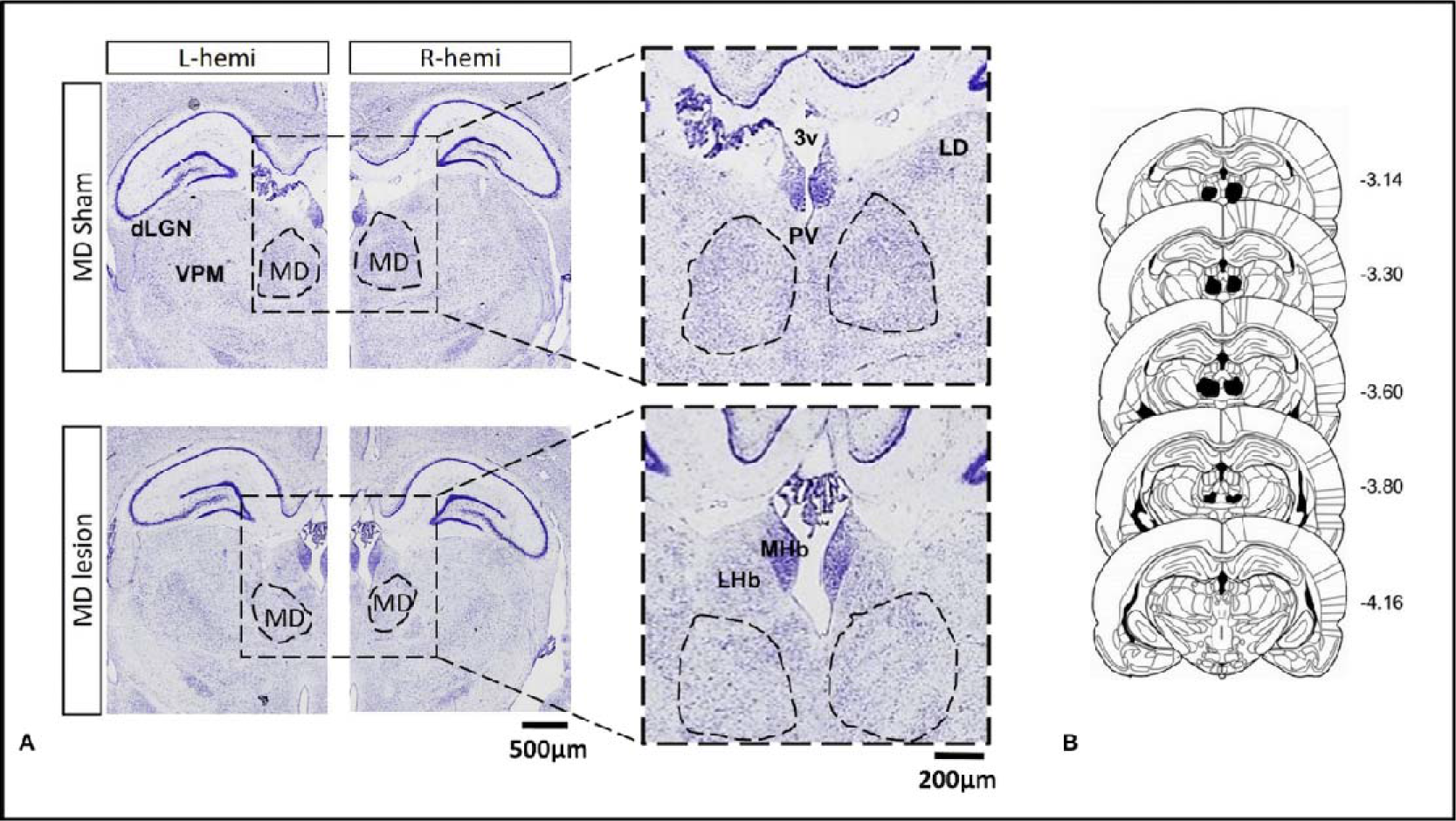
**A.** Photomicrographs of the thalamus in a MD Sham control rat (top) and a MD lesion rat (bottom). **B.** A series of coronal schematics throughout the mediodorsal thalamus showing the area of cell loss in the MD lesion group. Numbers refer to the distance from bregma (Paxinos and Watson, 2008). Abbreviations: 3v: third ventricle; dLGN: dorsal lateral geniculate nucleus; LHb: lateral habenula; LD: laterodorsal thalamus; MD: mediodorsal thalamus; MHb: medial habenula; PV: paraventricular nucleus; VPM: ventral posterior medial thalamus.

## 3.2 The MD and attentional set-shifting

Preoperatively, rats were pseudo-randomly assigned to either of two groups: MD lesion or MD Sham control, based on the number of trials needed to reach the learning criterion (six correct consecutive trials in nine trials) during the simple discrimination (SD) subtask. Preoperative data from one MD Sham control rat had to be discarded. An independent samples t-test confirmed there was no significant difference between the two groups during preoperative learning of the SD, *t*(22) = 0.095, *p* > 0.05, with rats from both groups requiring a similar number of trials to reach criterion, MD lesion (Mean = 9.67, S.D. = 2.44) and MD Sham control (Mean = 9.78, S.D.= 3.31). Overall trials to criterion during the preoperative training session are presented in **Fig. 2A**.

Postoperatively, a repeated measures ANOVA with Stage (7 subtasks: SD, CD, REV1 ID, REV2, ED, REV3) as the within-group factor and Lesion_type (MD lesion vs MD Sham control) as the between-group factor revealed a significant interaction, Stage × Lesion_type, *F*(6,138) = 63.65, *p* <0.001, a significant main effect of Stage, *F*(6,138) = 184.73, *p* <0.001, and a significant main effect of Lesion_type, *F*(1,23) = 20.22, *p* <0.001 (see **Fig. 2B**). To explore the interaction effect, *posthoc* comparisons of the simple main effects of Stage (corrected for multiple comparisons, *p* <0.007; alpha = 0.05 divided by 7 tests) were computed. For the SD subtask, MD lesion rats required more trials to criterion (M = 16.67, S.D. = 1.23) compared with MD Sham controls (M = 9.50; S.D. = 2.07), indicating that the rats with the MD lesion were slower to acquire the new sensory discrimination. An independent samples t-test confirmed this difference was significant, *t*(23) = 10.88, *p* <0.001. However, once the rats with MD lesion acquired the rewarded sensory dimension (to respond either to the digging media, or to the odor), they were unimpaired in subsequent subtasks that maintained the same rewarded sensory dimension, namely concurrent discrimination (CD): *t*(23) = 0.00, *p* =1.0, and intra-dimensional shift (ID): *t*(23) = 1.08, *p* < 0.291. MD lesion rats were also not impaired in the reversals (REV1 and REV2) of the reward contingencies associated with the learned exemplars completed after the CD subtask, REV1: *t*(23) = 2.31, *p* < 0.030, or after the ID subtask, REV2: *t*(23) = 1.34, *p* < 0.192, although, all rats were familiar with the reversal rule as they had completed the ID/ED task preoperatively (see Materials and Methods). Extended Data **Figure 1-1** shows the mean and 1^st^ and 3^rd^ quartile box plots with the smallest and largest numbers of postoperative trials to criterion (whiskers) for the individual rats during each of the subtasks.

**Figure 2.**
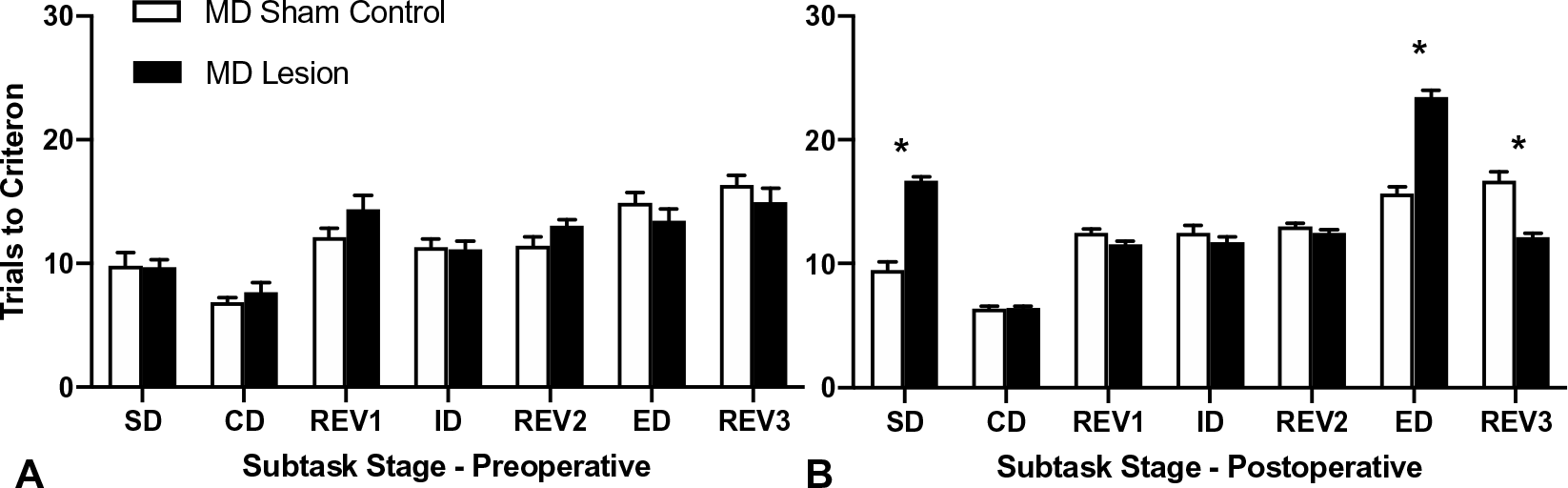
Mean (+SEM) trials to criterion data for the attentional set-shifting task. **(A)** Preoperative test session. **(B)** Postoperative test session. MD lesion rats were significantly slower to learn the SD and the ED subtasks compared to MD Sham controls (\**p* <0.001). MD Sham control rats, like MD lesion rats, needed more trials to learn the ED, as expected. However, the MD lesion rats required significantly fewer trials to criterion to learn REV3, the reversal occurring immediately after the ED, compared to the MD Sham controls (\**p* <0.001).

For the extra-dimensional (ED) subtask, the rats had to shift their attention to the previously irrelevant sensory dimension. Thus, given the rats had acquired the attentional set strategy, as expected, all rats required more trials to criterion to learn the ED subtask. MD Sham controls required (M = 15.70; S.D. = 1.64) more trials to learn the ED when compared to trials required to learn the ID (M = 12.5; S.D. = 1.84). Similarly, the MD lesion rats required more trials (M= 23.47, S.D. = 2.07) compared with trials to learn the ID (M = 11.73; S.D. = 1.67; see **Fig. 2B**). A repeated measures ANOVA computing the trials to criterion for these two subtasks (ID vs ED) showed a significant Stage x Lesion_type interaction, *F*(1,23) = 99.96, *p* <0.001, a significant effect of Stage, *F*(1,23) = 306.11, *p* <0.001, and a significant effect of Lesion_type, *F*(1,23) = 32.83, *p* <0.001 (see **Fig. 2B**). The interaction occurred as the MD lesion rats required significantly more trials to criterion to consistently implement a new choice strategy in order to learn which of two stimuli was rewarded from the previously ignored sensory dimension compared to the MD Sham controls, *t*(23) = 10.00, *p* <0.001.

Interestingly, for REV3, the reversal subtask that occurred after the ED shift, MD lesion rats required fewer trials to criterion (M = 12.13, S.D. = 1.25) compared with the MD Sham controls (M = 16.7; S.D. = 2.26; see **Fig. 2B**). An independent samples t-test confirmed this difference was significant, *t*(23) = 6.51, *p* <0.001, suggesting a facilitation in performance as a consequence of over-training experienced during the ED subtask (see discussion for interpretation).

The additional repeated measures ANOVA of Session (Preoperative vs Postoperative) shift cost (see **Fig. 3A**) between Lesion_type computed as the number of trials to criterion for the ED subtask minus the number of trials to criterion for the ID subtask for each session showed a significant Session x Lesion_type interaction, *F*(1,22) = 44.06, *p* <0.001, a significant effect of Sesion, *F*(1,22) = 38.27, *p* <0.001, and a significant effect of Lesion_type, *F*(1,22) = 15.46, *p* =0.001 (see **Fig. 3A**). The significant interaction occurred as there was a small shift cost for both groups during the preoperative test session. However, during the postoperative test session only the MD lesion rats showed a significant shift cost as a consequence of the task demands changing that required an adjustment to the previously well-established response strategy (*p* <0.001). By contrast, the MD Sham controls showed a similar shift cost to their preoperative performance.

**Figure 3.**
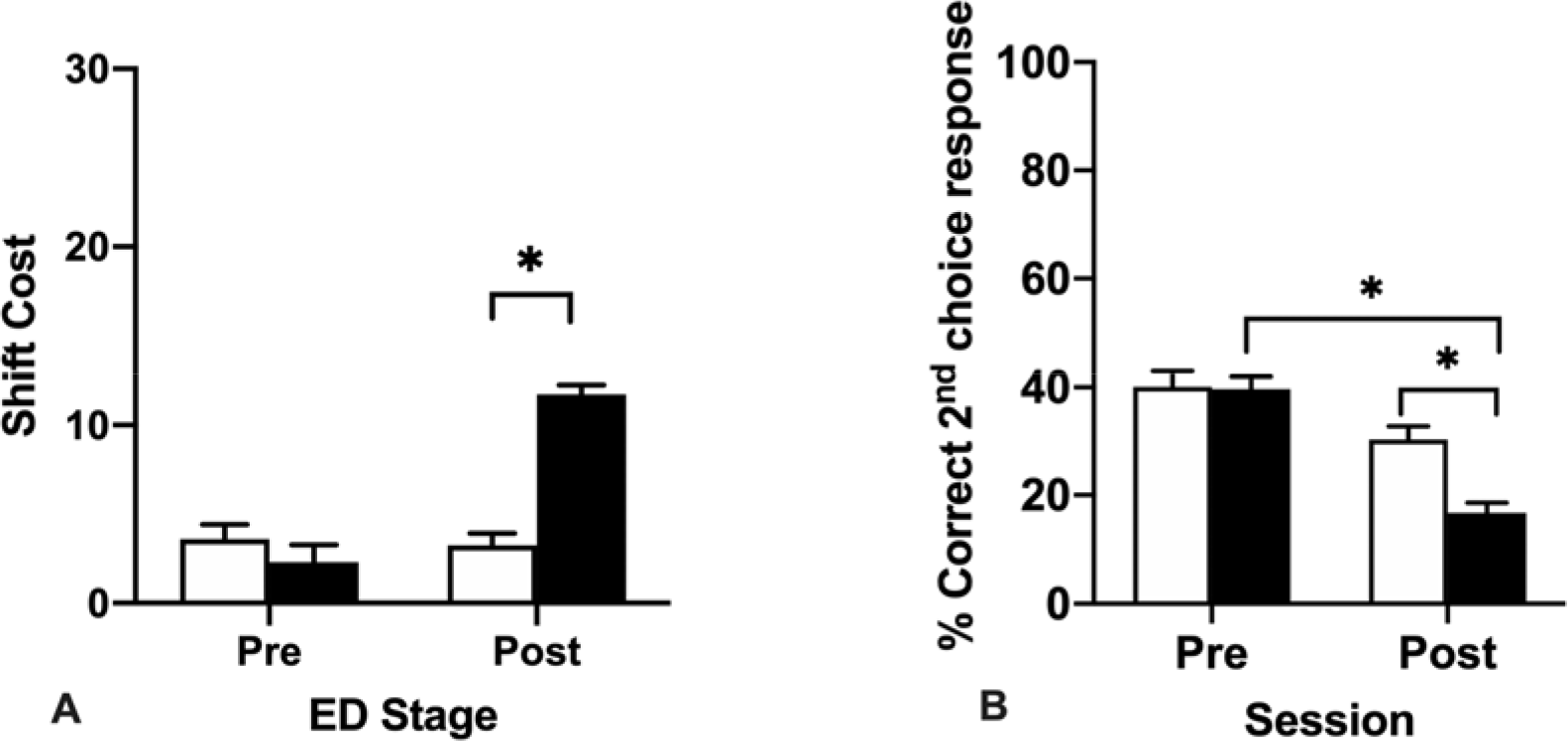
(**A**) Mean (+SEM) trials to criterion shift cost data for the ED stage for preoperative (Pre) and postoperative (Post) testing sessions. The shift cost is computed as the number of trials to criterion in the ED minus the number of trials to criterion in the ID. Both groups of rats showed a shift cost during the ED, when they had to learn to attend to the previously irrelevant dimension. However, the shift cost for the MD lesion rats in the postoperative test session was significantly different to the MD Sham controls. (**B)** Mean (+SEM) percent correct 2^nd^ choice within trial responses made during the preoperative and postoperative performance sessions of the attentional set-shifting task. MD lesion rats made fewer correct 2^nd^ choice within trial responses than MD Sham controls (* *p*’s = 0.001).

### 3.3 Within trial choice responses as a measure of rapid adaptability after MD lesion

Given the observed deficits in the MD lesion group, we investigated the behavioral responses made in the MD lesion or MD Sham control rats. Given the task design and a criterion of six consecutive correct responses before switching to the next stage, overall during testing, the rats do not complete many trials. Nevertheless, we could determined whether the rats rapidly adapted their choice responses in the here and now (within trial choice responses) as measured by the number of correct 2^nd^ choice within trial responses made, i.e. when the rat dug from the correct bowl only after visiting the incorrect bowl first but without digging in it, divided by the total number of correct responses (correct 1^st^ and 2^nd^ choice responses combined) for each session. During each subtask of the session, we recorded if the rat made a correct choice, either on the first attempt (‘1^st^ choice response’), or on the 2^nd^ attempt (‘2^nd^ choice within trial response’). We did not include error trials in this analysis.

A repeated measures ANOVA of total percent correct 2^nd^ choice within trial responses was conducted with Session (Preoperative vs Postoperative) as the repeated measure × Lesion_type revealed a significant interaction of Sesssion × Lesion_type, *F*(1,21) = 9.40, *p* = 0.006, a significant main effect of Session, *F*(1,21) = 57.06, *p* <0.001, and a significant main effect of Lesion_type, *F*(1,21) = 7.18, *p* = 0.014 (see **Fig. 3B**). To investigate the interaction effect, *posthoc* comparisons of the simple main effects showed that the MD lesion rats (M = 28.18, SD = 7.61) made fewer correct 2^nd^ choice within trial responses compared to the MD Sham controls (M = 35.23, SD = 8.46). This deficit suggests the MD lesion rats had a diminished ability to rapidly bind together their previous choice about what stimuli are worth sampling in order to update their choice response strategy within the trial, rather than a deficit in perseverative responding, or an inability to respond to negative feedback.

In addition, we analyzed the rats’ errors made reaching criterion during the ED subtask, as a consequence of whether they experienced a sensory dimension shift from odor to digging medium or digging medium to odor, which indicated that the increase in errors occurs irrespective of the direction of the attentional shift. The univariate ANOVA with Lesion_type and Sensory Shift as the between-subject factors and errors to criterion for the ED subtask as the dependent measure revealed a main effect of Lesion_type, *F*(1, 21) = 106.03, *p* < 0.001, but no main effect of Sensory Shift, *F*(1, 21) = 3.84, *p*= 0.063 and no interaction effect, *F*(1, 21) = 0.19, *p*=0.666.

### 3.4 Latency to dig changes

Latency to dig (in seconds) for errors were also computed for each rat in separate (preoperative (see **Fig. 4A**) and postoperative (see **Fig. 4B**)) analyses. Preoperatively, a repeated measures ANOVA with Stage (7 subtasks: SD, CD, REV1 ID, REV2, ED, REV3) as the within-group factor and Lesion-type (MD lesion vs MD Sham control) as the between-group factor revealed a significant main effect of Stage, *F*(1,21) = 23.99, *p* <0.001, but no effect of lesion, and no interaction, *F*’s <1.0. This result indicates there was no difference in response times between the pseudo-randomly assigned groups (MD lesion and MD Sham controls) preoperatively and that all rats responded quicker on errors trials as they moved through subsequent subtasks within the session.

However, a separate repeated measures ANOVA of latency to dig during postoperative error trials revealed a significant interaction of Stage × Lesion_type, *F*(6,132) = 26.81, *p* <0.001, a significant main effect of Stage, *F*(6,132) = 22.45, *p* <0.001, and a significant main effect of Lesion-type, *F*(1,22) = 57.15, *p* <0.001 (see **Fig. 4B**). To investigate the interaction effect, *posthoc* comparisons of the simple main effects (corrected for multiple comparisons) were computed. Postoperatively, MD lesion rats responded slower on error trials during the SD subtask, *t*(22) = 10.09, *p* <0.001, the CD subtask, *t*(22) = 7.65, *p* <0.001, and the REV1 subtask, *t*(22) = 4.44, *p* <0.001. However, for the final reversal subtask, REV3, completed after the ED shift, MD lesion rats were quicker on the error trials than MD Sham controls, *t*(22) = 3.53, *p* = 0.002. This result in combination with the reduced trials to criterion required to complete this subtask (see **Fig. 2A**) suggests that the MD lesion rats may have experienced overtraining due to the number of additional trials to criterion required to adapt their choice responding in the ED subtask (Dhawan et al., 2019). We did not include correct trials in this analysis as it is a forced choice task and both groups of rats were motivated to retrieve the Honey Nut Cheerios reward on each trial.

**Figure 4.**
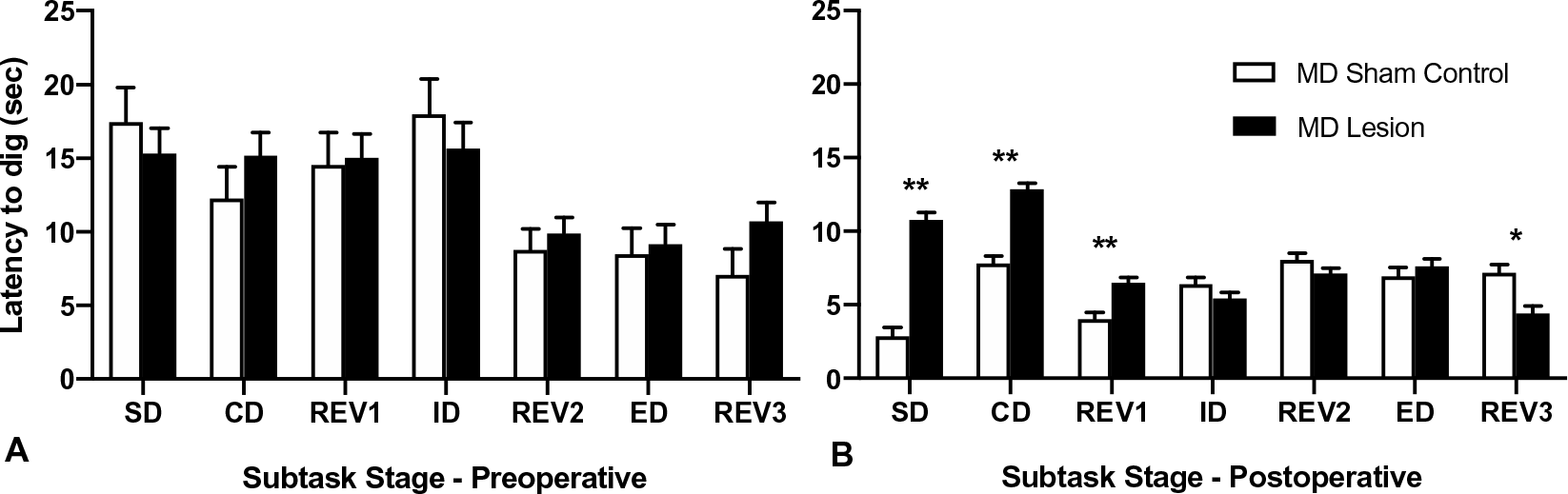
(**A**) Mean (+SEM) latency to dig (secs) for preoperative error trials for MD Sham controls and MD lesion rats during each subtask of the ID/ED attentional set shifting task. (**B**) Mean (+SEM) latency to dig (secs) for postoperative error trials for MD Sham controls and MD lesion rats during each subtask of the ID/ED attentional set shifting task (** *p* < 0.001; * *p* = 0.002).

## 4.0 Discussion

This set of experiments investigated the active influence of rat mediodorsal thalamus (MD) in cognitive flexibility. More specifically, we examined whether rats with excitotoxic lesions of the central and medial MD were impaired in performing an attentional set-shifting task that measures intra-dimensional (ID) and extra-dimensional (ED) shifts in attention to sensory discriminations (digging media or odors). Our new results indicate that rats with MD lesion required more trials to criterion to learn the optimal response strategy during the initial simple discrimination (SD). Although, once this stable response strategy has been learnt, MD lesion rats were able to attain an attentional set strategy as they acquired further two-choice discriminations involving the same sensory dimension during the CD and ID shifts, and during reversals (REV1 and REV2) of the reward contingencies associated to the exemplars. However, when the relevant sensory dimension was switched, for the ED subtask, the MD lesion rats were markedly impaired and required many more trials to update their choices using a new response strategy, as measured by the significant ED shift cost. Further, for REV3, the third reversal performed after the ED shift, the MD lesion rats showed facilitated learning, suggesting they had received ‘overtraining’ as a consequence of completing many more trials to reach criterion in the ED subtask. Overtraining in discrimination tasks can facilitate reversal learning (Reid, 1953) and has been studied for many decades in psychology (Lovejoy, 1966). The analyses of behavioral responses indicated that the MD lesion rats made significantly fewer correct 2^nd^ choice within trial responses suggesting that information transfer via the MD is critical for the rapid, within trial monitoring and updating of an optimal choice response. Interestingly, similar updating deficits in choice responding have been previously reported in rhesus macaques with magnocellular MD excitotoxic lesions (Chakraborty et al. 2016).

These behavioral data accord somewhat with previous studies conducted in several species that targeted equivalent frontal regions. For example, marmosets with lateral PFC lesions (Dias et al., 1996) or rats with medial PFC lesions (Birrell and Brown, 2000) are also impaired at the ED subtask, showing a similar ED shift cost as our MD lesion rats. Conversely though, lesions to these comparative PFC regions in marmosets and rats did not cause animals to need more trials to reach criterion during acquisition of the initial choice response strategy in the SD subtask, indicating frontal regions and dorsal thalamic regions contribute to different aspects of updating cognitive flexibility. We attribute the differences between our learning results and the study of Birrell and Brown (2000) to differences in the investigated brain areas. In Birrell and Brown (2000) the lesions were centered on prelimbic cortex and included damage to infralimbic cortex. In the rat, the MD projects to medial prelimbic cortex, orbitofrontal cortex and cingulate (Cg1) cortex. A large body of research indicates that separate rat prefrontal cortex subregions differentially contribute to learning, memory, and other cognitive functions (for review, see Dalley, Cardinal and Robbins, 2004; Uylings, Groenewegen and Kolb, 2003).

Interestingly though, in relation to the dorsal thalamus, while rats with anterior thalamic nuclei (ATN) lesions are not impaired in the ED shift, they are impaired at acquiring the SD subtask (Wright et al., 2015). However, in contrast to our MD lesion rats, rats with ATN lesions continued to be impaired during the CD subtask and subsequent ID shifts (Wright et al., 2015) indicating that they never properly learnt the discrimination rule, and suggesting an intact ATN (a brain structure adjacent to the MD but with markedly different cortico-thalamo-cortical connectivity) is important for supporting the formation of an attentional set strategy.

Conversely, with more trials, our rats with MD lesions were able to acquire the optimal choice response strategy during the SD subtask and apply this optimal rule during subsequent subtasks, indicating that they had required the correct response rule, but they were markedly impaired at rapidly adapting this now well-established choice response strategy when they had to attend to the previously irrelevant sensory dimension during the ED shift. The demands on cognitive flexibility, response inhibition, and adapting behavioral responses after negative feedback are increased when the animals must make an ED shift, assuming they have acquired the ID attentional set strategy in the first place (Dhwana et al., 2019). The increased effort involved in adapting to these changes in attention is measured by a shift cost, and it is expected that more trials to criterion are required during learning the ED than during learning the ID shift. While the ID/ED shift cost was higher for both groups (Fig. 2B), it was significantly increased for the MD lesion rats compared to the MD Sham controls. As indicated, increased shift costs have also been observed in marmosets and rats with lesions to the medial PFC (Dias et al., 1996; Birrell and Brown, 2000). Surprisingly though, rats with damage to the nucleus reuniens (Re), another thalamic structure located near the MD, which interconnects the medial PFC and hippocampus (Hoover and Vertes, 2007; Hoover and Vertes, 2011; Vertes et al., 2015) do not cause deficits in the ED subtask or produce a significant ED shift cost. Instead, rats with NRe damage are impaired at acquiring the attentional set strategy, similar to rats with ATN lesions (Linley et al., 2016). Thus, our MD thalamus results are unique.

Other researchers using mice have proposed the MD supports the frontal cortex to sustain intra-cortical attentional control without transferring categorical information about a particular task rule (Schmitt et al., 2017), while others have shown medial PFC-MD projections are important for behavioral flexibility but not task engagement (Marton et al., 2018; Nakayama et al., 2018). Further, the MD has been shown to have a critical role in maintaining the balance between excitation and inhibition in dorsomedial PFC via its influence on interneurons (Anastasiades et al., 2021; Ferguson and Gao, 2018) and pyramidal neurons (Collins et al., 2018; Delevich et al., 2015). Our results are supportive of these circuit level interactions, although our deficits indicate the influence of MD on frontal circuits is specific to certain aspects of the attentional set-shifting task, e.g. linked to the ED shift and learning the optimal choice response strategy, but not to the ID shift or completing reversals. However, our study involved permanent MD lesions, which are likely to have caused some adaptations across the network, while the other studies likely involve short-term changes as temporary inactivation of the MD was employed. Nevertheless, it may be proposed that after MD perturbations, while the animal is still able to detect a change in the task demands (i.e. they are responsive to negative feedback), they are impaired at rapidly adapting their behavior and coordinating correct choice responses within a trial after a change to the already established choice response strategy (Block et al., 2007; Hunt and Aggleton, 1998; Chakraborty et al., 2016). Interestingly, in both the rodent studies, the deficits were attributed to increased perseverative responding. Others have also observed increased perseverative responding after MD perturbations (Ferguson and Gao, 2018). In contrast, our current rodent work and previous work in rhesus macaques has shown an intact MDmc is necessary for rapid reward guided learning of complex discriminations (Mitchell et al., 2007a; Chakraborty et al., 2016; Chakraborty et al., 2019), although the deficits linked to learning in monkeys are attributed to increased response switching (Mitchell et al., 2007a; Chakraborty et al., 2016). These congruent results across species after MD damage suggest that cortical information transfer via the MD is particularly important when rapid, within a trial changes in choice response strategies linked to establishing a new rule are required.

In addition, while the selectivity of the deficits after MD lesion are in some ways comparative to medial PFC perturbations on attentional set-shifting, i.e. both have a significant ED shift cost, our MD lesion rats also took more trials to criterion during the simple discrimination (SD) subtask. Other studies also confirm MD lesion rats are impaired in acquisition of an initial task rule (Hunt and Aggleton, 1998). Other rodent studies causing temporary perturbations to the MD also show impaired discrimination learning (Ferguson and Gao, 2018; Courtiol et al., 2019). Thus these results from across species reinforce the notion that MD is not simply mimicking the PFC (DeNicola et al., 2020; Miller et al., 2017; Mitchell et al., 2007; Mitchell and Gaffan 2008; Mitchell et al., 2014). Consequently the influence of other cortico-cortical interactions that also indirectly transfer information to the cortex via the MD and other thalamic structures, e.g. those PFC interactions with limbic structures in the temporal lobes and sensory association areas are also critical during learning sensory discriminations (Banjeree et al., 2020; Bueno-Junior et al., 2018; Chakraborty et al., 2019; Floresco and Grace, 2003; Mitchell, 2015; Pelekanos et al., 2020).

For the reversal subtasks (REV1 and REV2), surprisingly, MD lesion rats were unimpaired during reversals in the reward contingencies. In contrast, rats or monkeys with OFC perturbations are impaired in the reversal subtasks of the ID/ED task but conversely do not show an ED shift cost (Dias et al., 1997; McAlonan and Brown, 2003; Chase et al., 2012). MD is reciprocally connected to the OFC (Groenwegen, 1988; Preuss, 1995; Ray and Price, 1992). Thus this double dissociation in deficits between these two reciprocally interconnected brain regions needs to be reconciled. First, it must be noted that the rats had experienced all of the subtasks (and thus were familiar with the concept of reversal learning and that it can appear in the task structure) during their preoperative test session, which may have reduced or eliminated the reversal learning deficit (Costa et al., 2015; Jang et al., 2015). Interestingly though, rats with OFC lesions completing the ID/ED task also had preoperative exposure to the concept of reversal learning but as indicated, they were impaired at reversal learning during their postoperative testing (McAlonan and Brown, 2003). Thus, this contrasting evidence suggests the MD and OFC are computing different aspects of cognitive flexibility. Further, for the MD, while previous evidence of reversal learning deficits are reported for some tasks after MD perturbations (Block et al., 2007; Chudasama et al., 2001; Ferguson and Gao, 2018; Hunt and Aggleton, 1998; Ostlund and Balliene, 2008; Parnaudeau et al., 2013, 2015), the evidence is mixed (Beracochea et al., 1989; Alcaraz et al., 2018; Fresno et al., 2019). These mixed effects may be the consequence of differences in task structure and the way that the reversal is introduced. Other factors, including the extent of the networks disrupted by the MD perturbations that spread to adjacent thalamic structures may also be a factor (Mitchell and Chakraborty, 2013; Wolff and Vann, 2019). Indeed, one recent study has identified that a nearby thalamic structure, the submedius thalamus may contribute a role in reversal learning, instead of the MD (Fresno et al., 2019). Thus, as already indicated, the MD is not simply mimicking the behavioral effects observed after PFC damage.

Consequently, our current results and this above evidence highlights that the information transfer between OFC and MD is not always critical for reversal learning per se. Instead, at least in monkeys, OFC-MDmc interactions are critical for adaptive, value based decision-making (Browning et al., 2015; Izquierdo and Murray, 2010; Mitchell et al., 2007b), while in rodent studies, the OFC-striatal part of the cortico-striatal-thalamic neural circuits have been implicating in reversal learning (Gremel and Costa, 2013; Izqueirdo et al., 2017; Nakayama et al 2018; Yang et al., 2020; Yin et al., 2005) or OFC-submedius thalamus circuits (Fresno et al., 2019). Consequently other OFC-striatal-thalamic and OFC-thalamic interactions are potentially more involved in supporting reversal learning. In addition, thalamic inputs from the intralaminar thalamic nuclei and motor thalamus to the striatum contribute a selective role in inhibitory control and behavioral flexibility (e.g. Saund et al., 2017).

Intriguingly, our additional analyses used to investigate the types of behavioral deficits occurring showed that our MD lesion rats made fewer correct 2^nd^ choice within trial responses. Given that MD lesion rats required more trials to criterion to learn the SD and ED subtasks, this change in behavioral responses suggests that without the MD thalamus, our rats could not rapidly adapt their start of trial choice responses as the trial was progressing (within trial) after the changed task demands. Interestingly though, this reduced correct 2^nd^ choice within trial responding does not suggest the rats adopted perseverative responding (they would have potentially been impaired in reversal learning if they had) or were not learning about the stimulus and associated rewards. We can conclude this because, after acquiring the ED shift, the MD lesion rats required fewer trials to criterion to learn the final reversal, REV3. This facilitation of learning during the final reversal subtask suggests that the MD lesion rats had experienced ‘overtraining’ on all of the stimulus features associated with each sensory dimension as they required more trials to learn the ED subtask (Dhawan et al., 2019). As Pearce and Mackintosh (2010) indicate, until learning is fully consolidated, all stimulus features continue to be attended to and this seemed to be so for our rats, even with the MD lesion. Thus, we propose, in accord with others, that through experiencing these extra trials during the ED subtask, the MD lesion group may have developed a greater understanding of the rewarded and unrewarded stimulus dimensions, thus increasing the salience of the predictive cues and reducing/eliminating the number of factors that can led to an error. As indicated above, it is well established that overtraining rats during discrimination learning can eliminate any reversal learning deficits (Reid, 1953; Lovejoy, 1966; Nilsson et al., 2015 for review).

There may be other explanations for the SD and ED impairments observed in our MD lesion rats. For example, with the ID/ ED task, it is important to establish that the animals understand the relevant sensory dimension that is rewarded across the related subtasks within the session. Fortunately, in our animals, this transfer of knowledge was evident as the MD lesion rats showed reliable learning in the subtasks, CD, REV1, ID and REV2, that required the implementation of choice responses (follow the same rule) to the same sensory dimension as the SD to receive reward. Furthermore, both Sham controls and MD lesion rats showed similar numbers of trials to criterion during the REV1 and REV2 subtask, which indicates that they did not favor one feature of the stimulus dimension more than the other or that they were insensitive to negative feedback. In addition, we only used male rats in this current study so the results might not transfer to female rats. Finally, the MD is a subcortical node in the olfactory neural circuitry that also includes the OFC (Courtiol et al., 2015; Veldhuizen et al., 2020). However, evidence collected from rodents with MD perturbations or humans with strokes affecting the MD indicates that changes to the MD do not impair olfactory discriminations (Courtiol et al., 2019; Tham et al., 2011). Moreover, our MD lesion rats showed similar levels of olfactory discriminations as the MD Sham controls (see Material and Methods for further details).

Finally, the diffuse influence of the MD thalamocortical inputs to several frontal cortex structures (Alcaraz et al., 2016; Kuramoto et al., 2017; Mitchell, 2015; Wolff and Vann, 2019) following MD lesion supports previous findings implicating medial PFC-MD connectivity in flexible behaviors (Marton et al., 2018; Nakayama et al., 2018). Furthermore, the MD along with the OFC provides additional evidence for the contribution of this area in supporting online, ‘here and now’ reward guided learning and decision-making (Gardner et al., 2017; Gardner et al., 2019; Schoenbaum et al., 2011; Wallis, 2007). Intact OFC connectivity is essential to the animals’ ability to perform reward-guided learning in order that other interconnected neural networks, likely including cortico-cortical, cortico-striatal, and cortico-thalamo-cortical connectivity, can determine the optimal choice responses and implement the appropriate actions (Browning et al 2015; Chakraborty et al 2016; Nakayama et al 2018; Perry et al., 2021; Rushworth et al., 2011).

The ID/ ED task is analogous to the Wisconsin Card Sorting Test (WCST) in humans. In healthy humans performing the WCST during neuroimaging, the MD is activated during responding to negative feedback after the choice has been executed (Monchi et al., 2001). Unfortunately, thus far, humans with stroke damage in the MD have cognitive deficits that are clinically very poorly defined (Pergola et al., 2018). However, people diagnosed with Alzheimer’s disease (AD), Parkinson’s disease (PD), or schizophrenia show impaired attentional set-shifting performance (e.g. Barch et al., 2009; Monchi et al., 2004; Owen et al, 1993). Neuroimaging and post-mortem studies show marked changes in the MD and/ or ATN in these diseases (Hornberger et al., 2012; Ouhaz et al., 2018; Pergola et al., 2015; Perry et al., 2018). Our current evidence advocates for studies investigating cortico-thalamocortical transfer of information in people diagnosed with AD combined with more frontal pathology, or PD, or schizophrenia.

To conclude, excitotoxic damage to the rodent central and medial MD selectively increased the trials to criterion on the ED subtask, producing a shift cost. This selective performance deficit is similar to monkeys with lateral PFC inactivation (Dias et al., 1996; Dias et al., 1997) and rats with medial PFC lesions (Birrell and Brown, 2000). Further, this dissociable deficit after MD lesion contrasts with monkeys or rats with perturbations to the OFC, who are selectively impaired on reversal learning, but not on ED shifts (Chase et al., 2012; Dias et al., 1997; McAlonan and Brown, 2003). As evidenced, the frontal cortex is critically involved in value based decision-making and reward guided learning (Miller, 2000; Rushworth et al. 2011; Wallis, 2007). However, cortico-thalamo-cortical connections also contribute a role (Browning et al. 2015; Chakraborty et al., 2016; Izqueirdo and Murray, 2010; Mitchell et al., 2014; Pelekanos et al., 2020). In rodents, the MD and the medial PFC together appear crucial for binding reward information and behavior (Bradfield et al., 2013; Corbit et al., 2003), as inhibition of dorsomedial PFC-projecting MD neurons results in rats having difficulties with tracking changes in action-outcome contingencies (Alcaraz et al., 2018). In the current study, we show the rat medial and central MD is critical for rapidly updating an optimal choice response strategy. That is, when the MD is damaged, there is a diminished ability to rapidly learn (within a trial) a choice response strategy, as well as update a well-established choice response strategy as task demands change within a session.

However, instead of increased perseverative errors as observed in other rodent studies with MD lesions, our MD lesion rats made less correct 2^nd^ choice within trial responses, suggesting they were unable to rapidly monitor and update within trial changes of mind decisions when they encountered the unrewarded stimuli on a given trial.

## Acknowledgements

We are very grateful for support received from Professor Verity Brown and Dr David Tait, who taught ZO to perform the ID/ED attentional set-shifting task during an extended visit to their research lab at St Andrews University.

---

**Extended Data Figure 1-1.**
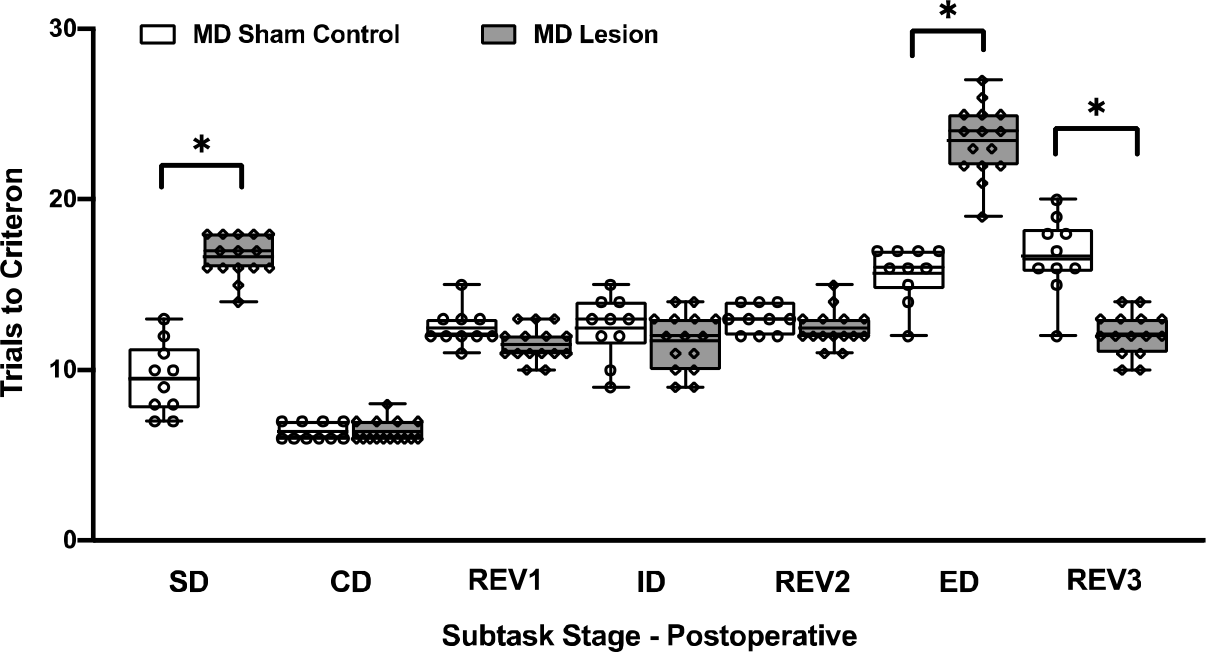
Boxplots showing mean trials to criterion for individual rats during the postoperative sessions of the attentional set-shifting task. MD lesion rats took significantly more trials to learn the SD and the ED subtasks compared to MD Sham controls (\**p* <0.001). MD Sham control rats also needed more trials to learn the ED, as expected. The MD lesion rats required significantly fewer trials to learn REV3, the reversal occurring immediately after the ED, compared to the MD Sham controls (\**p* <0.001). Box shows 1^st^ and 3^rd^ quartile and whiskers are the minimum and maximum trials to criterion for individual rats.

